# TDAExplore: quantitative image analysis through topology-based machine learning

**DOI:** 10.1101/2021.06.13.448249

**Authors:** Parker Edwards, Kristen Skruber, Nikola Milićević, James B. Heidings, Tracy-Ann Read, Peter Bubenik, Eric A. Vitriol

## Abstract

Machine learning has greatly expanded the ability to classify images. However, many machine learning classifiers require thousands of images for training and lack quantitative descriptors of how images were grouped. We overcome these limitations with a machine learning approach based on topological data analysis, where a data set of 20-30 images is sufficient to accurately train the classifier. Our method quantifies differences between groups and identifies subcellular regions with the largest dissimilarities.

---

Microscopy images contain an incredible amount of complex information. The development of machine learning has substantially accelerated the ability to identify and extract relevant features, as well as to classify images into groups [1,2]. These methods, such as convolutional neural networks, perform extraordinarily well in segmentation and classification [3,4]. However, identifying how machine learning pipelines interpret training data to determine which features to extract is challenging.

Here, we present an image analysis pipeline based on two topological data analysis (TDA) methods, persistent homology [5] and persistence landscapes [6]. TDA is a mathematical method that uses algebraic topology to quantify the shape of data [7]. After training, our TDA-based classifier identifies regions whose shape features most strongly characterize each class of image. Using multiple data sets, we demonstrate that this method achieves high per-image classification accuracies with minimal training, is robust to hyperparameter changes, and requires only modest computational resources. Subsequently, we show this approach can recognize a wide variety of subcellular structures and extract biologically meaningful data.

To extract topological information, images were first masked by automatic intensity thresholding to reduce pixel values outside of the cell to zero. Images were then divided into uniform radius patches (Fig. 1A: i). A subset of high intensity pixels were selected per patch (Fig. 1A: ii). Neighboring points were progressively connected if they are within an increasing distance of each other. (Fig. 1A: iii-vi). This sequence of simplicial complexes was then used to generate a persistence landscape (Fig. 1A: viii, B), which encodes sets of birth-death pairs (Fig. 1A: vii) generated from the appearance and disappearance of specific homology features [10], including connected components and the holes within them.

**Figure 1.**
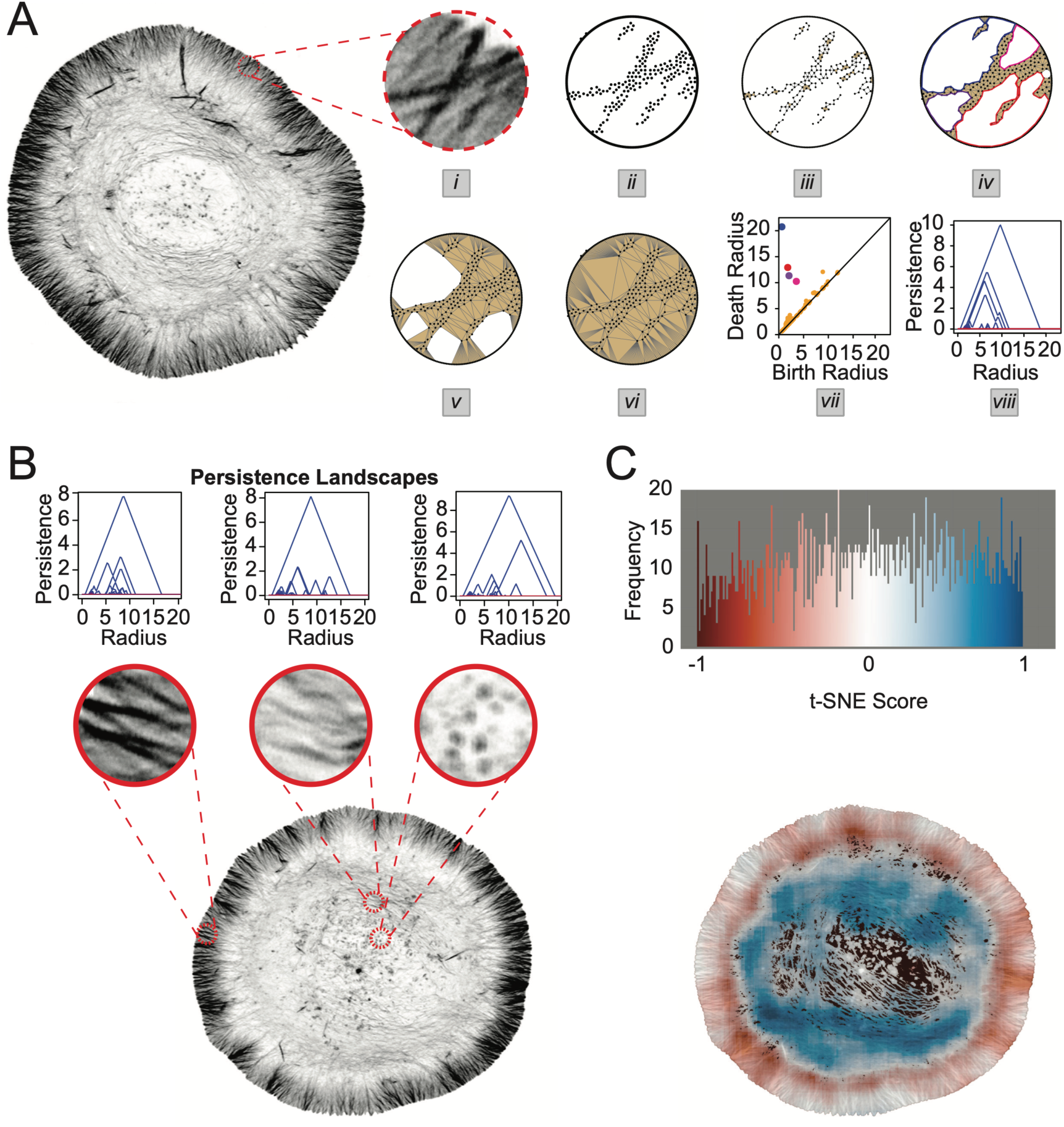
Persistent homology-based image analysis pipeline to extract topological features. **(A)** Computation of persistence landscape from selected image patch. (i) Representative image patch of 75 pixel radius (ii) Points are sampled from spatial coordinates of fluorescence signal (iii-vi) Sequence of simplicial complexes generated by connecting neighboring points within a distance that increases from (iii) to (vi). (iv) Four cycles that represent persistent homology classes in degree one are colored. (v) The pink cycle has been filled in. (vi) All cycles have been filled in. (vii) Persistence diagram plotting the birth radius and death radius of each of the persistent homology classes in degree one. Colored points correspond to the colored cycles in (iv). (vii) Persistence landscape gives a vector encoding of the persistent homology. **(B)** Persistence landscapes generated from each selected patch in the input image. Three representative patches and their landscapes are shown. **(C)** Histogram of t-SNE scores generated from persistence landscapes. Colorized pixels representing t-SNE scores overlaid on the original image as a mask.

To interpret the intrinsic characteristics of the resulting high dimensional vectors, t-distributed Stochastic Neighbor Embedding (t-SNE) [11] was used to reduce image summaries to a single score. The scores were subsequently scaled to range from −1 to 1 and a color gradient was mapped to the values. These values were then mapped to the pixels of the corresponding uniform radius patch so that the result can be visualized on the input image as a mask (Fig. 1C). These masks show that this method extracts physiologically relevant features of the cytoskeleton by demarcating the large actin superstructure known as the lamellipodia from the cell interior (Fig. 1C).

We then turned to machine learning methods to enhance the classification power of the TDA descriptors from each data set. Two main hyperparameters were chosen to run the pipeline: the radius in pixels of the subsampled patches and the number of patches sampled per image (see Supp. Equation 1). A uniform selection of pixel radius 75 and patch ratio 2 was made for all data sets based on biological considerations, namely the size in pixels of relevant subcellular structures.

Our study was restricted to linear support vector regression (SVR) classifiers to ensure that our results depended on the quality of the TDA descriptors and not on the strength of the machine learning method. To train a 2-class image classification model, persistence landscapes were generated from image patches. Support vector machines defined a hyperplane that optimizes the separation between training data with different class labels [13] and subsequently assigned a score to each patch and persistence landscape. The classifier identified patches with score < 0 as one class, while those with score > 0 are in the other. Score values were scaled to range from −2 to 2. To classify an image, the patch scores for the image were averaged, and a class was assigned corresponding to whether the average score was < 0 or > 0. We used 5-fold cross validation to assess classifier performance (Supp. Fig. 1). The images in a data set were separated into 5 approximately equal data groupings or “folds” with training and testing at an 80/20 ratio (Supp Fig. 1 A). The number of correctly identified images out of the total number was visualized as a confusion matrix (Supp. Fig. 1 D). To assess performance for the entire pipeline, we performed 5-fold cross validation 50 times for each data set. (Supp. Table 1).

This combination of TDA and machine learning (TDAExplore) was then used to detect changes to the architecture of the actin cytoskeleton after induction of both genetic and chemical perturbations to major actin assembly factors (Supp. Table 2). We found TDAExplore was able to robustly identify which cells were challenged with perturbations to the actin cytoskeleton (Fig. 2, Supplementary Fig. 2). This included using a small molecule inhibitor (CK666) against the actin nucleator Arp2/3 (Fig. 2 A), CRISPR/Cas9 knockout of the gene for the actin monomer binding protein profilin 1 (Fig. 2 B), shRNA knockdown of the actin filament severing protein cofilin 1 (Supp. Fig. 2 A, D) and the monomer binding protein Thymosin β-4 (Supp. Fig. 2 B, E), and mislocalization-based inhibition of the Mena/VASP actin polymerases (Supp. Fig. 2 C, F). To ensure that TDAExplore was not overfitting input data by detecting non-relevant features, controls from all actin datasets were randomized and tested against each other. TDAExplore was not able to separate features of control datasets, demonstrating that bona fide changes to actin architecture are required to make distinctions between data sets (Supp. Fig. 3 E).

**Figure 2.**
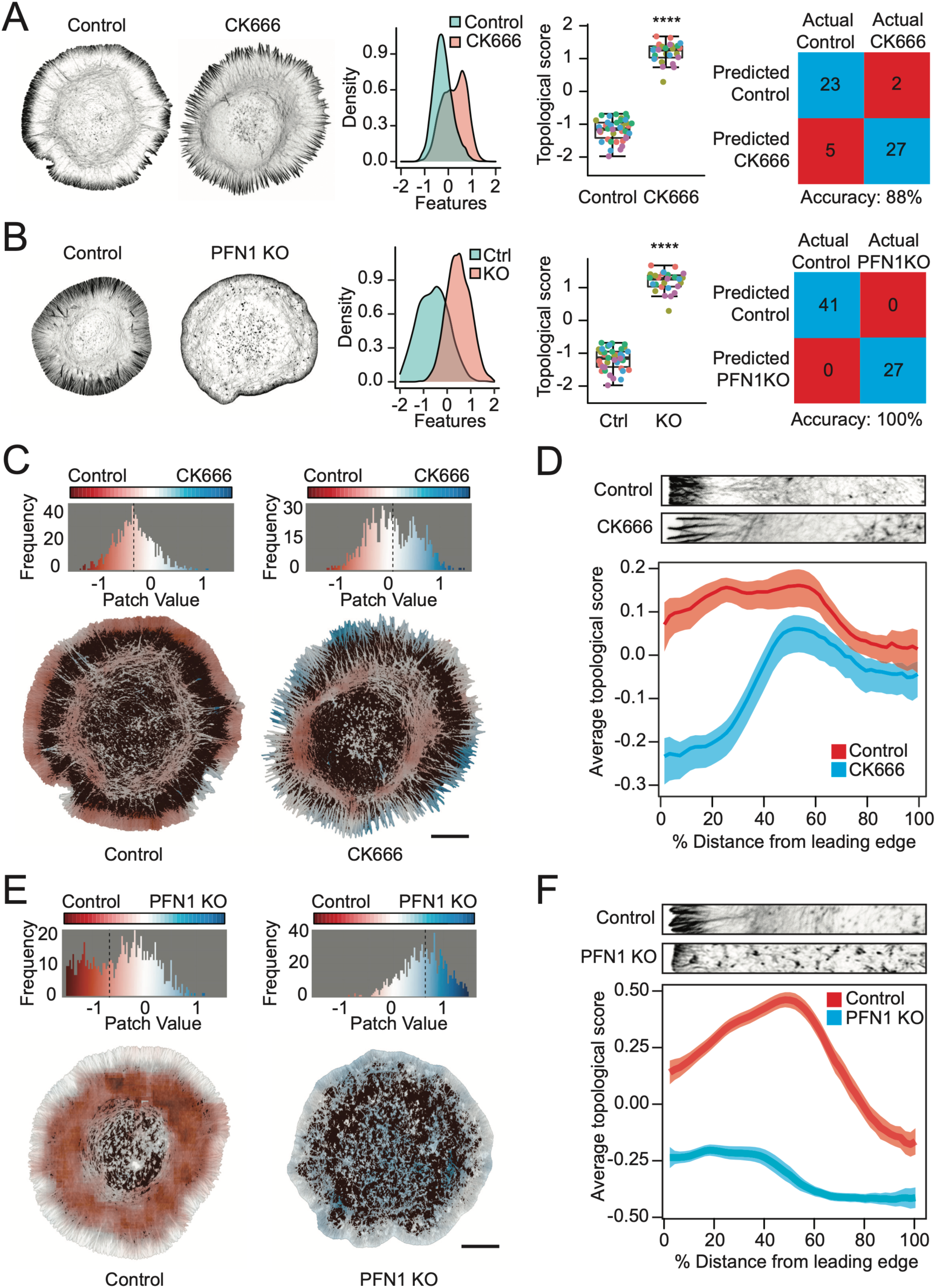
TDAExplore analysis of the actin cytoskeleton. **(A)** Performance evaluation for classification of cells treated with the Arp2/3 inhibitor CK-666 or its inactive control CK-689. N= 25, 32 for CK-689 and CK-666, respectively. From left to right, distribution of patch features, topological score per cell, and confusion matrix displaying classification summaries. Topological scores are generated through five separate rounds of testing, rounds are designated by color. **(B)** Performance evaluation for classification of control and PFN1 KO cells, n= 41, 27, respectively. From left to right, distribution of patch features, topological score per cell, and confusion matrix displaying classification summaries. Topological scores are generated through five separate rounds of testing, rounds are designated by color. **(C)** Distribution of patch values after treatment with CK689 and CK666 where CK689 patches are values <0 and colored red, while CK666 is classified as patch values >0 and colored blue. White represents intermediate values. Below is the computed feature masks of patch values for representative images from CK689 and CK666 treated cells. Scale bar represents 10µm. **(D)** Average topological score based on distance from the leading edge to the cell center. Transparent bands depict 95% confidence intervals. Representative cell regions from the leading edge to center are shown above for comparison. **(E)** Distribution of patch values for control and PFN1 KO cells where control patches are values <0 and colored red, while PFN1 KO is classified as patch values >0 and colored blue. White represents intermediate values. Below is the computed feature masks of patch values for representative images from control and PFN1 KO cells. Scale bar represents 10µm. **(F)** Average topological score based on distance from the leading edge to the cell center. Transparent bands depict 95% confidence intervals. Representative cell regions from the leading edge to center are shown above for comparison. Graph depicting the difference between average topological score as related to distance from the leading edge.

To test whether this method is suited for detecting other structures, we altered mitochondrial morphology using the oxidative uncoupler carbonyl cyanide m-chlorophenylhydrazone. The classifier still performed well with an 81% accuracy between groups (Supp Fig 4 B). We also used TDAExplore on other published data sets (summarized in Supp. Table 3) which were previously classified with machine learning. TDAExplore performed comparably well to these methods, including on data from low-resolution or non-fluorescent images (Supp. Table 3, Supp. Fig. 6 G).

Next, we used TDAExplore to extract spatial information at the subcellular level. In a process similar to our t-SNE mask generation, patch values generated from SVR on persistence landscapes were mapped back to their pixels of origin so that topological scores could be visualized on the input image as a mask. This provides a visualization of the spatial effect of each perturbation with red colored pixels classified as CK689 and blue pixels mapped to CK666 (Fig. 2 C). To test whether the process of mask production is sensitive to changes from random sampling, a PFN1 KO classifier was trained five times and masks were generated for each classification run. Masks were consistent across training sessions, indicating that random sampling did not have a significant effect on mask generation (Supp. Fig. 3 F).

To further quantify spatial components of the topological score, the center of each cell was defined and patch values were assessed based on distance to the center (Fig. 2 D, F). Arp2/3 inactivation with the small molecule inhibitor CK666 causes a striking rearrangement of actin architecture at the cell boundary without changing total polymerized actin levels [8,9]. Our feature mapping recapitulated these results and showed that the average topological score for CK666-treated cells deviated most strongly from their controls near the cell boundary (Fig. 2 C, D). Knockout of PFN1 (Fig. 2 E, F) and depletion of cofilin (Supp. Fig. 5 C, D) also showed differential effects based on subcellular location. The largest topological score difference between control and PFN1 KO cells was approximately 50% of the distance from the boundary to the cell center, while the largest difference in topological scoring between cofilin KD cells and control was at the cell boundary. The deviation in topological scoring did not map strongly to specific subcellular locations for Tβ4 KD cells, although there was a slightly larger difference at the cell interior than at the edge (Supp. Fig. 5 A, B). 100% image accuracy in the profilin 1 knockout (PFN1 KO) dataset (Fig. 2 B) may indicate both the strong effects and penetrance of the perturbation. The relatively weaker training performance of both knockdown/genetic depletion of Tβ4 and cofilin (84% and 91%, respectively) (Supp. Fig 2) may either indicate that their depletion induces fewer effects on actin architecture or inform on penetrance of the depletion itself (i.e. the possibility that not all cells were depleted to the same extent).

One advantage of TDAExplore is its high performance using small datasets (on average 60 images per set), made feasible by assessment at the patch level. The possibility that changes to dataset size could alter image accuracy was tested by reducing high performing datasets in half. When the number of images was reduced from 68 to 34, 100% accuracy for the PFN1 KO classifier was maintained (Supp. Fig. 3 D). Because these data sets were generated from the same cell line, controls for the genetic knockout, depletion, inactivation and small molecule inhibition datasets should have indistinguishable features. When controls from all experiments were tested, their patch scores were not statistically different from each other (Supp. Fig. 3 E).

To understand how accessible TDAExplore would be to users with different computational resources, we collected usage statistics with our PFN1 KO data set (Supp. Fig. 7 A, B) for multiple hyperparameter selections. Our results indicate that with 8 processors, 5-fold cross validation with patch radius 75 pixels and patch ratio 2 would take less than 13 minutes and 5 GB of memory. The pipeline can utilize additional processors to further decrease computation time.

In summary, TDAExplore combines TDA and machine learning for segmentation and feature extraction of high resolution images. It is unbiased, relying on minimal user input for image exploration, segmentation, and classification. It is able to differentiate between multiple types of perturbations over a wide range of subjects without overfitting. Importantly, it also visualizes features at the sub-image level. Thus, TDAExplore should be well suited to extract novel quantitative information from imaging data in a wide variety of applications.

## Methods

### Cell culture

All cells used in this study are Cath.-a-differentiated (CAD) cells (originally purchased from Sigma-Aldrich). CAD cells were cultured in DMEM/F12 medium (Gibco) supplemented with 8% fetal calf serum, 1% L-Glutamine, and 1% penicillin-streptomycin. Prior to imaging, cells were plated on coverslips coated with 10 μg/mL Laminin (Sigma). No new cell lines were generated in this study and cell line generation is detailed elsewhere. For the generation of Cofilin knockdown refer to Vitriol et al., 2013 (26); For details on the generation of Thymosin β-4 knockdown refer to Lee et al., 2013 (27); For PFN1 knockout, Arp2/3 inactivation, Mena/VASP sequestration refer to Skruber et al. 2020 [8].

### Immunofluorescence

For visualization of actin filaments, cells were fixed with 4% electron microscopy grade paraformaldehyde (Electron Microscopy Sciences) for 10 min at RT and then permeabilized for 3 minutes with 1% Triton-X. Cells were then washed three times with PBS and actin filaments were stained with Alexa Fluor 488 phalloidin or Alexa Fluor 568 phalloidin (diluted 1:100, Life Technologies) for 30 min at room temperature in immunofluorescence staining buffer. For visualization of mitochondria, cells were cells were fixed with 4% electron microscopy grade paraformaldehyde (Electron Microscopy Sciences) for 10 min at RT and then permeabilized for 5 minutes with 0.1% Tween-20. Cells were washed three times with PBS and incubated with an antibody to TOM-20 (rabbit, 1:500, CST) at 4°C overnight. Cells were then washed 2X in PBS and incubated with (goat anti-rabbit Alexa Fluor 568, 1:1000) for 2 hrs and then washed 3X in PBS before mounting with Prolong Diamond (Life Technologies).

### Microscopy

Images were acquired with a Nikon A1R+ laser scanning confocal microscope with a GaAsP multi-detector unit. All confocal imaging was performed with an Apo TIRF 60X 1.49 NA objective. Deconvolution-based super-resolution confocal microscopy [10] was performed by using zoom settings higher than the Nyquist criteria, resulting in oversampled pixels (0.03 μm). Confocal z-stacks were created and then deconvolved with Nikon Elements software using the Landweber algorithm (15 iterations, with spherical aberration correction) to create images with approximately 150 nm resolution [11]. For input into the analysis pipeline, cell boundaries were identified by thresholding and all signal outside of the cell was reduced to zero. Maximum intensity z-projections were created from the bottom of the z-stack excluding the top of the cell.

### Third party datasets

We used image datasets BBBC 013v1, 014v1, 015v1, and 016v1 from the Broad Bioimage Benchmark Collection [12] and Binucleate and Liver Gender datasets from the Image Informatics and Computational Biology Unit (IICBU) Biological Image Repository [13] to validate our TDA approach. BBBC data are images of human osteosarcoma cell (U2OS) from high throughput microscopy assays. Only the GFP channel images were analyzed. If the data contained multiple images from the same well, only one image per well was used for the analysis to mitigate batch effects. Subsets of data with only two classes of phenotypes were selected using the procedure described by Uhlmann et al. [14]. IICBU data contain images from D. melanogaster cells and murine liver tissue. Only the red color channel was analyzed for murine liver tissue images.

### Extracting persistence landscapes from images

Gray scale images were first summarized by sampling patches and computing persistence landscapes as described in Fig. 1. This method optimizes computational demands while maintaining analysis quality. Patches were selected from each image by repeatedly sampling patch centers from a probability distribution constructed on the image’s pixels. The probability distribution was biased towards higher intensity pixels to prioritize patches centered in non-background regions. The selection also integrated a geometric criterion: newly sampled centers were removed if they were within 2 pixels’ distance of a previously sampled pixel. This method was an alternative to grid-based approaches used for convolutional neural networks, and minimizes patch overlap while focusing on regions of interest. Because it involved a probability distribution, this step was not deterministic.

Each selected patch was converted to a set of 2D points by taking the coordinates of the top 2.5% of pixels by intensity while observing a similar geometric criterion. This resulted in point samples which captured the most intense regions in patches while avoiding over-sampling. Each patch sample was augmented by adding a circle of points along the patch’s circular boundary. This enabled efficient computation of persistent local homology, which captures features like straight lines in addition to holes [15]. Alpha complex persistence diagrams for 0 and 1 degree homology were computed using the R package TDA [16] which wraps the software GUDHI [17] for such computations. Persistence landscapes were computed using the R package tda-tools [18] which wraps the Persistence Landscapes Toolbox [19]. The first 50 landscapes were retained for each degree of homology, and each was discretized using 201 values for a total 10050 dimensions per homology degree in the final output vectors.

### Support vector regression on patch landscapes

Two class (control vs experimental group) support vector regression models were trained on a diverse range of data sets. For each data set, images were used to generate patches and corresponding persistence landscapes. Persistence landscapes were labeled based on treatment.

Every model training session consisted of first splitting the set of images into a testing set (approximately 20% of the data) and a training set (approximately 80% of the data). The labeled patch landscapes from training images were used to train a linear SVR model (L2 regularized and L2 loss) via the R wrapper for Liblinear [20]. The cost parameter C was selected using the package’s built-in heuristic on the training landscapes. Per-class cost weights were used to penalize misclassification of training points in the smaller class more heavily. Accuracy on patches was assessed by comparing labels assigned to the testing data by the model to the actual labels.

The trained SVR model assigned a score to each patch landscape. These scores were averaged for each image to assign aggregate scores to all training and testing images. A second SVR was trained similarly to the first, but with the training image scores and labels. Image accuracy was assessed by comparing predicted image labels to actual labels for the testing set.

### Assessing model performance

A single round of model assessment on a data set consisted of first generating persistence landscapes from the data set, then performing 5-fold cross validation by training and testing both patch and image level SVR models from the persistence landscapes. The relatively small size of the data sets induced substantial variance in accuracy across folds. The total number of correctly classified testing images across folds was reported, rather than average testing accuracy across folds.

Since patch selection had a non-deterministic component, performance on each data set was assessed by conducting 50 independent rounds of model assessment with the same hyperparameters and recording the average accuracy across the 5 cross validation folds for each round. Supp. Table 1 reports summary statistics for these accuracies on each data set. Standard deviations for image classification accuracy were uniformly less than 5% across all data sets, indicating low variance induced by random sampling.

### Hyperparameter selection

The radius of patches (in pixels) and the number of patches sampled per image were hyperparameters in the persistence landscape generation procedure. The number of patches per image was selected as a “patch ratio” obtained by dividing the total number of pixels in an input image by the summed area of all patches to be sampled from an image.

The standard practice of tuning hyperparameters by splitting data sets into hyperparameter training, model training, and model testing sets was infeasible given the small number of images in the data sets. Instead, a uniform patch radius of 75 pixels and patch ratio of 2 was used for all models. The radius was initially selected based on the size of biologically relevant features in the PFN1 data set, while the patch ratio of 2 was selected arbitrarily.

To verify that this choice of hyperparameters did not produce misleading results, model performance using a range of hyperparameter values was assessed. For each data set, two single rounds of model assessment were performed for each choice of hyperparameters with patch radii taking values 30, 60, 90, 120, 150, and 180 and patch ratios taking values 0.33, 0.67, 1.00, 1.33, 1.67, and 2.00. In total there were 36 combinations of hyperparameters, each assessed twice, for 72 single rounds per data set. Altering the patch radius or ratio did not significantly affect image accuracy for our CK666 data set (Supp. Fig. 3 B). More generally, we also found that classification accuracy with patch radius 75 pixels and patch ratio 2 was not outlying for any data set (Supp. Fig. 3 C).

### Image masks from patch scores

The SVR models and t-SNE dimensionality reduction allowed the assignment of scores to image patches based on the patches’ persistence landscapes. To visualize these scores on the original images, a modified version of kernel convolution was used. Patches and persistence landscapes were computed for an image, and each pixel in the mask was assigned the average score of patches containing the pixel.

For t-SNE based scoring, mask pixels were then scaled to values between −1 and 1. 2000 patches with radius 75 were sampled in total.

Masks based on SVR classification were generated for all images in each data set. Mask pixels were scaled to between −1 and 1 across the entire data set. The number of patches and pixel radius were the same as those used for training the model.

### Line summaries from image masks

To further explore the spatial distribution of topological differences identified by the SVR models, image masks were converted to line summaries for actin-labelled data sets. The centroid of each image and the distance of a mask’s pixels from its centroid were computed. Distances were scaled between 0 and 1, and the average mask intensity for pixels with distances in bins of width 5% (0 to 5%, 5 to 10%, etc.) recorded. The resulting line summaries contained 20 scores, one for each bin. For each image class the average line summary and confidence bands containing 95% of scores were computed. Line summaries were visualized by linearly interpolating between the 20 data points.

### Computational performance assessment

Computational performance was assessed by conducting 5-fold cross validation rounds on the PFN1 data set for a range of hyperparameter values. Selected patch radii values were 25, 50, 75, 100, and 125. Patch ratio values were 0.4, 0.8, 1.2, 1.6, and 2. In total there were 25 combinations of hyperparameters, each assessed once.

Computations were performed using the University of Florida Research Computing’s “HiPerGator” cluster. Parallelization was utilized with 24 computation threads (reported as “CPU’s”) on a single Intel Gold 6142 2.6GHz processor. Resource usage statistics were collected using the monitoring software REMORA [21]. Supplementary Figure 4 records the results. Resource demands scale with the total number of persistence landscapes computed, which in turn are directly proportional to the patch ratio divided by the patch radius squared (see Supp. Equation 1).

## Software and data availability

TDAExplore is available as a command line program at: https://github.com/P-Edwards/TDAExplore-ML. It is also available as an R package for programmatic access at: https://github.com/P-Edwards/TDAExplore. Data used to generate this manuscript’s figures and tables and scripts for image analysis are available at: https://github.com/P-Edwards/TDAExplore-Examples

## Conflict of Interest

The authors declare no conflict of interest.

## Acknowledgements

Computations were performed using the University of Florida Research Computing’s supercomputer “HiPerGator”. This project was supported by a National Institutes of Health (NIH) Pathway to Independence Award (R00 NS087104) and an NIH Maximizing Investigators’ Research Award for Early Stage Investigators (R35 GM137959) to E.A.V. This research was partially supported by the Southeast Center for Mathematics and Biology, an NSF-Simons Research Center for Mathematics of Complex Biological Systems, under National Science Foundation Grant No. DMS-1764406 and Simons Foundation Grant No. 594594. This material is based upon work supported by, or in part by, the Army Research Laboratory and the Army Research Office under contract/grant number W911NF-18-1-0307.

## Notes

### Competing Interest Statement

The authors have declared no competing interest.

