## Supplementary Information for "TDAExplore: quantitative image analysis through topology-based machine learning"

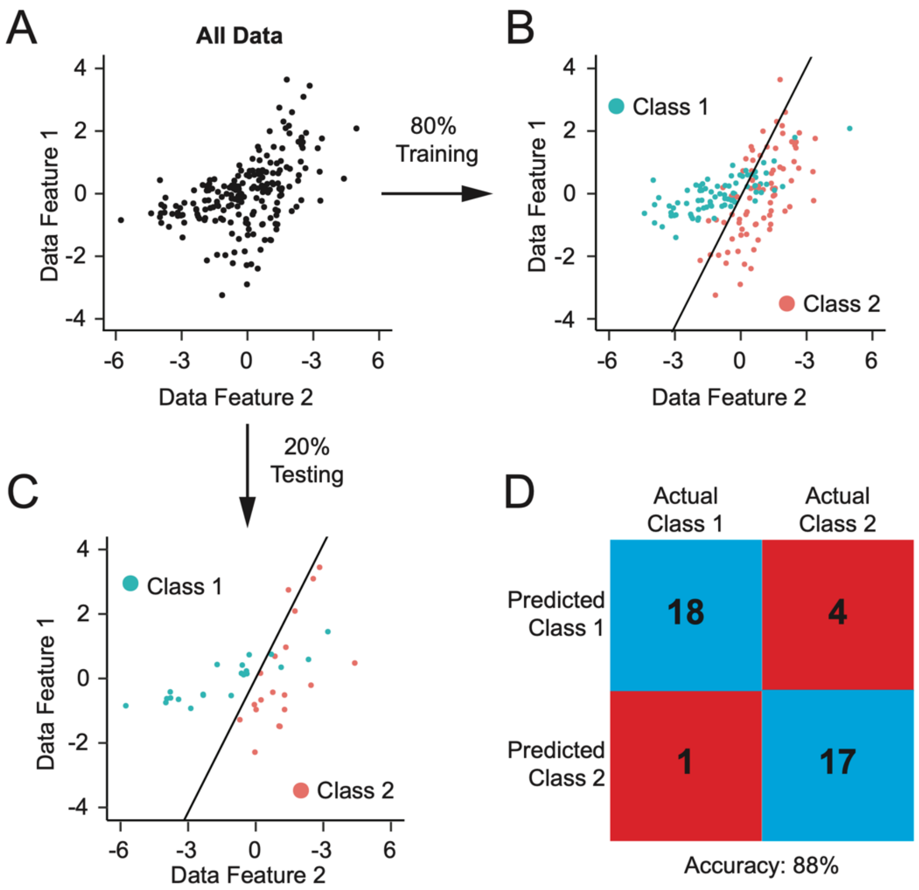


**Supplementary Figure 1. Example of TDAExplore classification pipeline**. **(A)** Data set separation into 5 approximately equal data groupings or "folds" with training and testing at an 80/20 ratio. Data are subjected to 5 rounds of training **(B)** or testing **(C)**, the black line indicates the hyperplane or classification boundary. **(D)** Example confusion matrix for visualization of image testing accuracy indicating number of correctly identified images out of the total number.

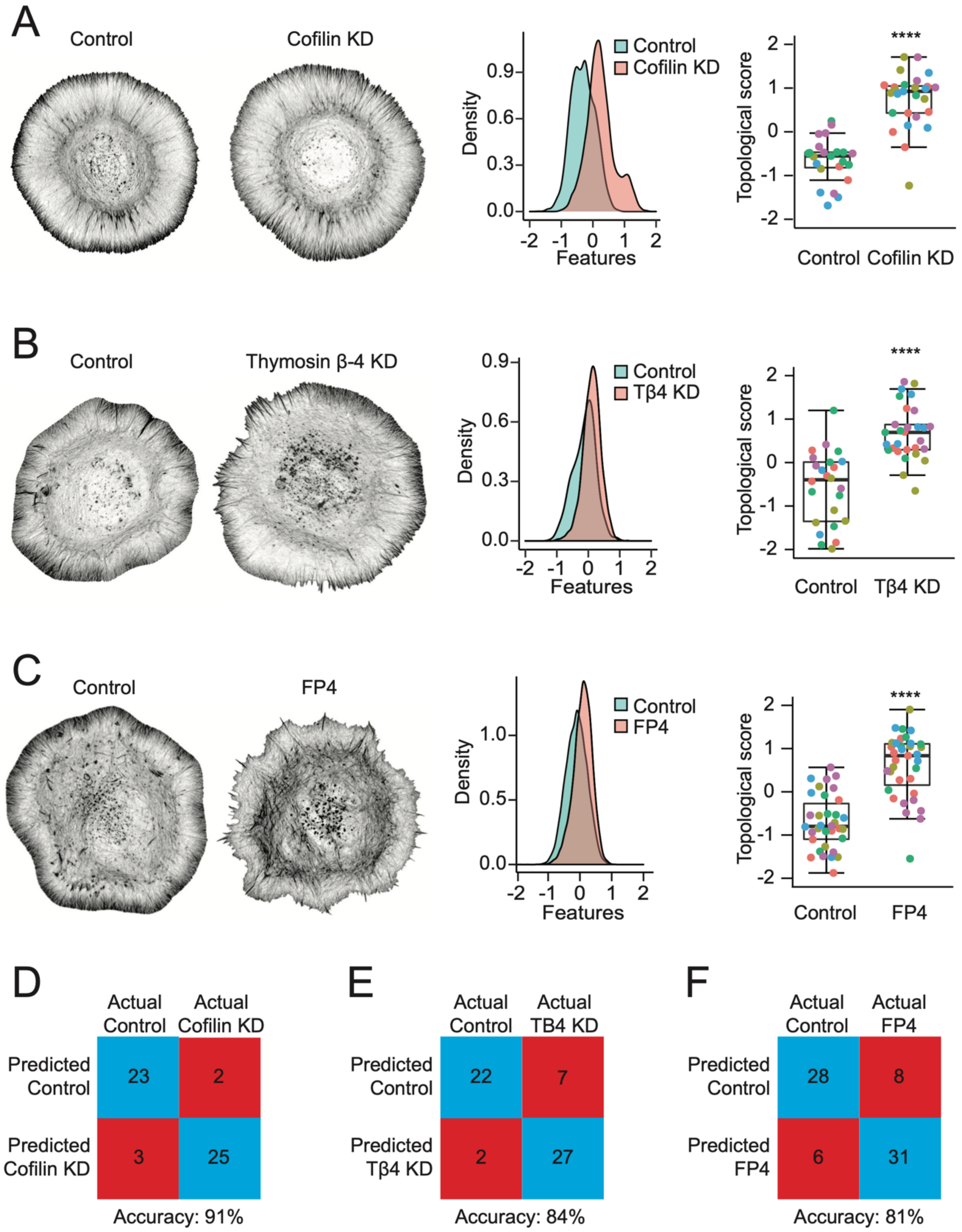


**Supplementary Figure 2**. **TDAExplore classification of additional perturbations to the actin cytoskeleton. (A)** Performance evaluation for classification of control knockdown and cofilin knockdown cells, n=25, 28, respectively. **(B)** Performance evaluation for classification of control knockdown and thymosin β-4 knockdown (Tβ4 KD), n=29 for both. **(C)** Performance evaluation for classification of cells treated with the Mena/VASP inhibitor FP4-Mito or its inactive control AP4-Mito. N= 36 for both. **(D-F)** Confusion matrices summarizing cofilin knockdown (D), Tβ4 KD (E), and FP4-Mito datasets (F).

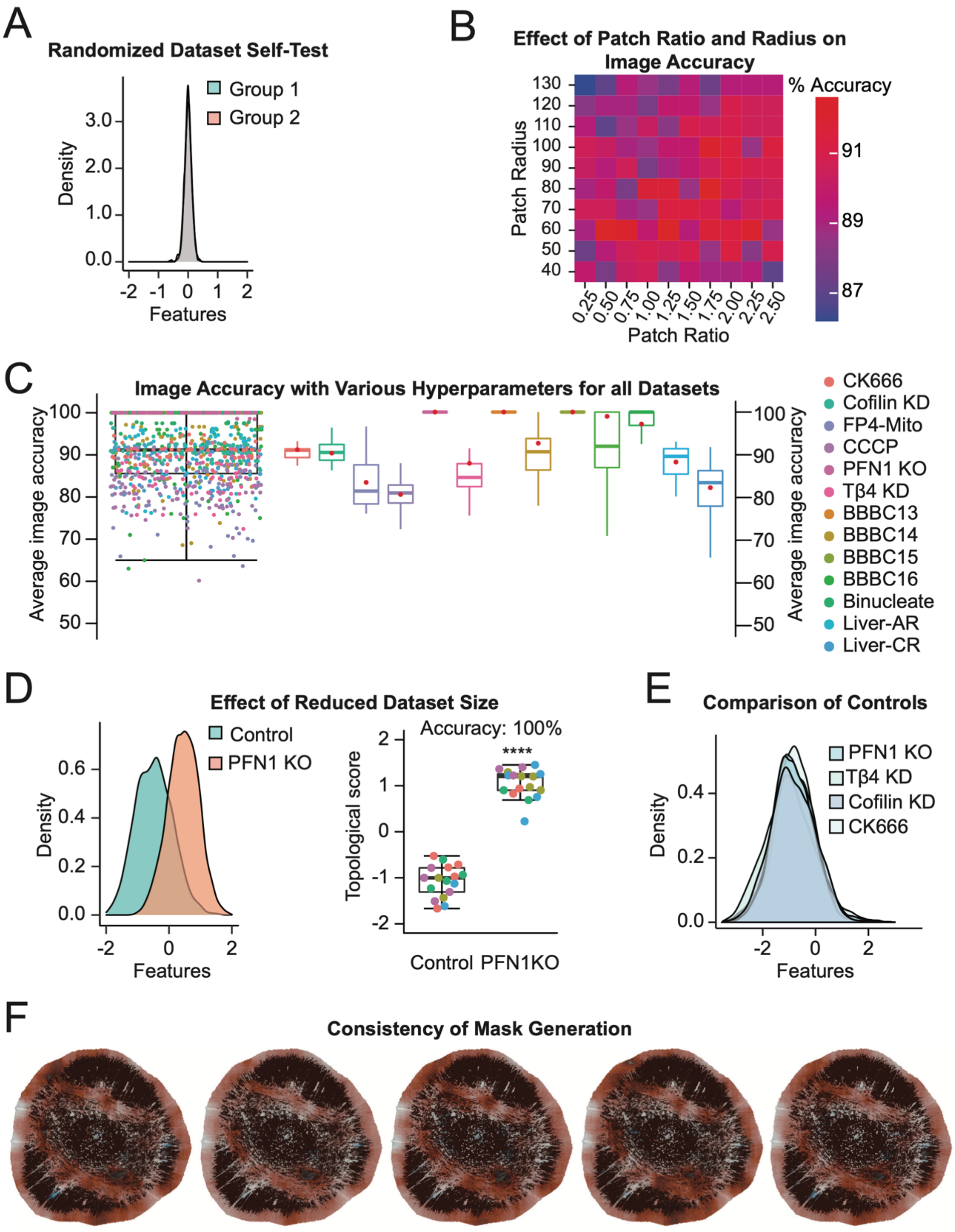


**Supplementary Figure 3. Input criteria do not strongly influence TDAExplore classifier performance. (A)** Randomization and testing of control data for PFN1 KO dataset. Images were arbitrarily split into groups of 21 and 20 and tested against each other. **(B)** Average image accuracy for the CK666 dataset after alterations to both patch radius and ratio. Accuracies are for whole image classification. **(C)** Average image accuracy for all datasets tested with TDAExplore and model assessment on each dataset for multiple choices of hyperparameters, for a total of 72 models per dataset. Accuracies are for whole image classification. Red dots indicate average classification accuracy for 50 models with patch radius 75 pixels and patch ratio 2. **(D)** Performance of the PFN1 KO dataset with only half the total images (N=34). **(E)** Patch feature distribution for controls of PFN1KO, Tβ4 KD, Cofilin KD, and CK666 datasets. N= 27, 29, 28, and 25, respectively.

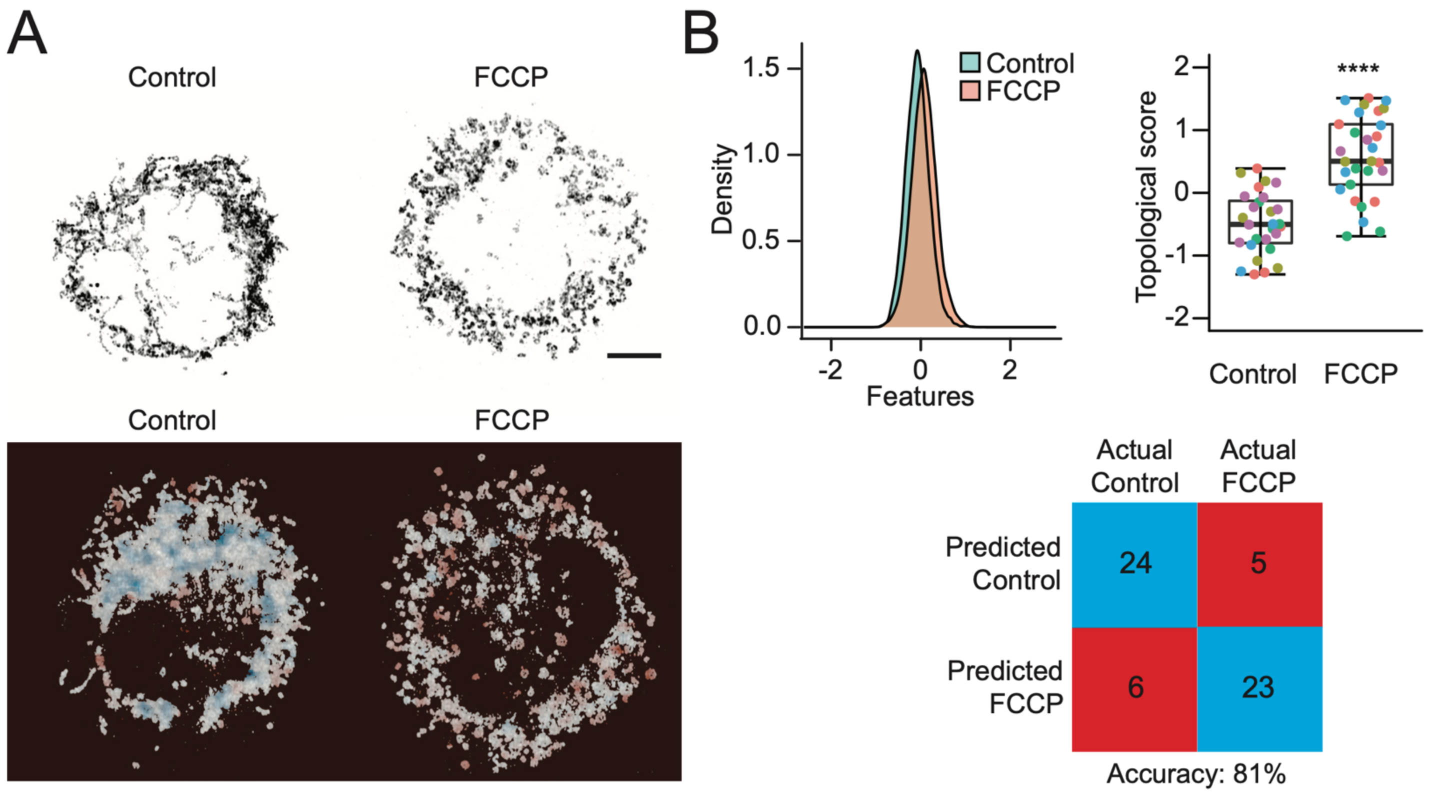


**Supplementary Figure 4. TDAExplore analysis of mitochondrial morphology. (A)** Representative images (top) and computed feature masks (bottom) of patch values for representative images from DMSO and CCCP treated cells. Scale bar represents 10μm. Performance evaluation for classification of cells treated with the oxidative uncoupler carbonyl cyanide m-chlorophenylhydrazone (CCCP) versus a DMSO control. N= 29 for both DMSO and CCCP. **(B)** From left to right, distribution of patch features, topological score per cell, and confusion matrix displaying classification summaries. Topological scores are generated through five separate rounds of testing, rounds are designated by color.

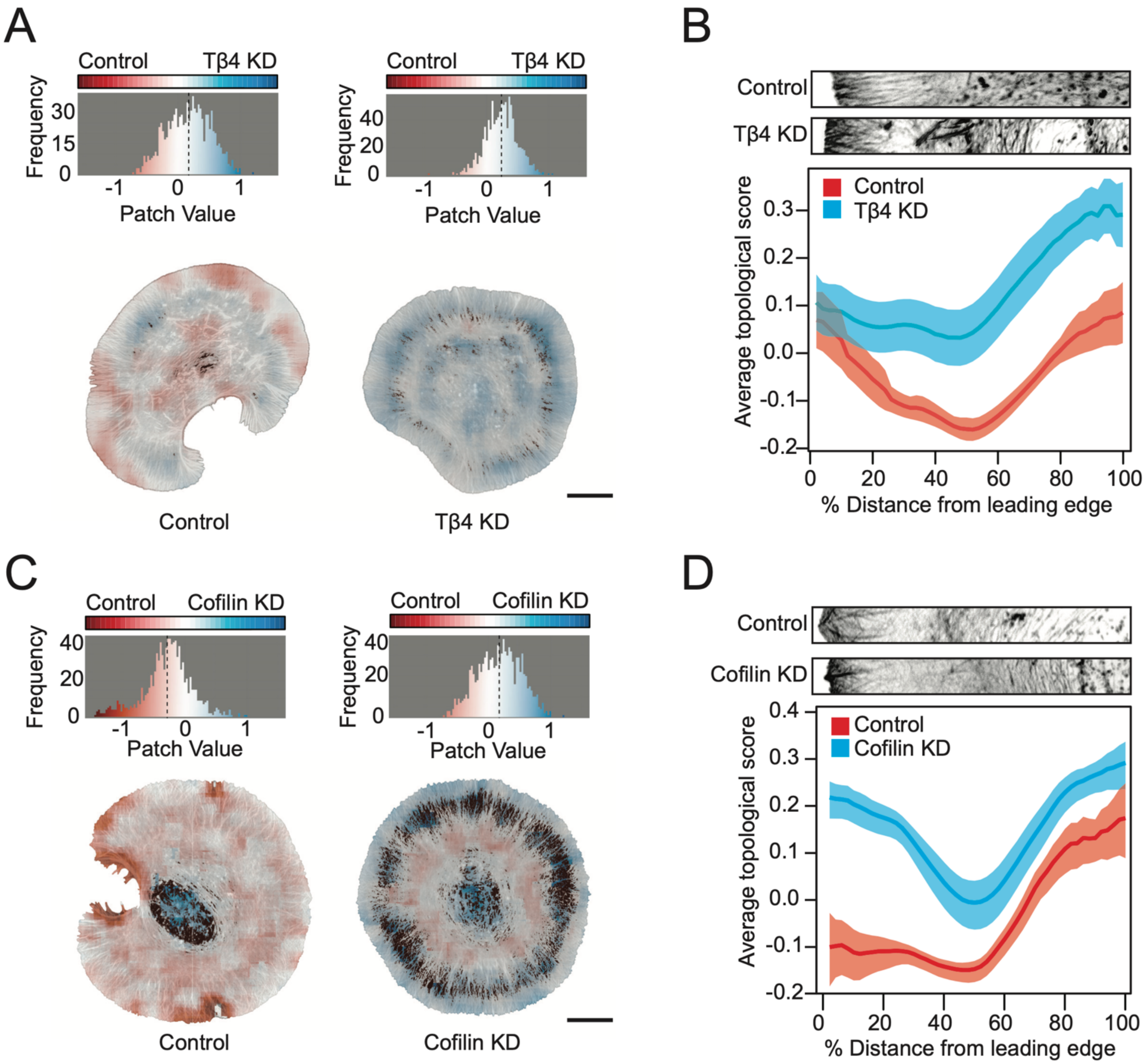


**Supplementary Figure 5. TDAExplore spatial analysis of additional perturbations to the actin cytoskeleton. (A)** Distribution of patch values after treatment for control and Tβ4 KD cells where control patches are values <0 and colored red, while Tβ4 KD is classified as patch values >0 and colored blue. White represents intermediate values. Computed feature masks (bottom) of patch values for representative images from control and Tβ4 KD cells. Scale bar represents 10μm. **(B)** Average topological score based on distance from the leading edge to the cell center. Transparent bands depict 95% confidence intervals. Representative cell regions from the leading edge to center are shown above for comparison. **(C)** Distribution of patch values after treatment for control and cofilin KD cells where control patches are values <0 and colored red, while cofilin KD is classified as patch values >0 and colored blue. White represents intermediate values. Computed feature masks (bottom) of patch values for representative images from control and cofilin KD cells. Scale bar represents 10μm. **(D)** Average topological score based on distance from the leading edge to the cell center. Transparent bands depict 95% confidence intervals. Representative cell regions from the leading edge to center are shown above for comparison.

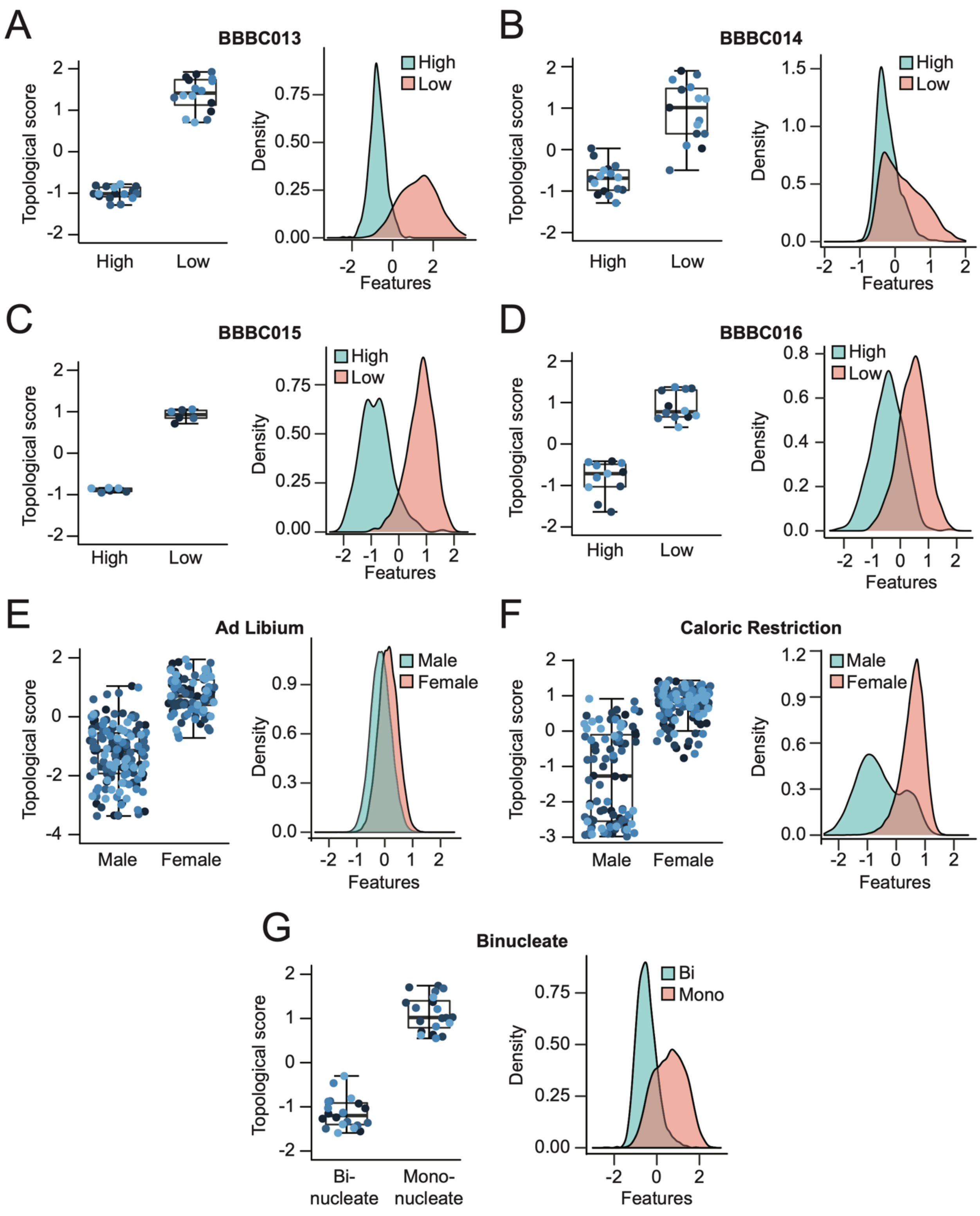


**Supplementary Figure 6**. **Performance evaluation of third party datasets using TDAExplore.** Performance evaluation for classification of the following: **(A)** BBBC dataset 013, n=16 images for both high and low. **(B)** BBBC dataset 014, n=16 images for both high and low. **(C)** BBBC dataset 015, n=6 for both high and low. **(D)** BBBC dataset 016, n=12 for both high and low. **(E)** IICBU dataset "Ad Libium". N=115, 150 for images from liver samples from female and male mice, respectively. **(F)** IICBU dataset "Caloric Restriction". N=153, 150 for images from liver samples from female and male mice, respectively. **(G)** IICBU dataset "Binucleate". N=20,21 images from binucleated cells and mononucleated cells, respectively.

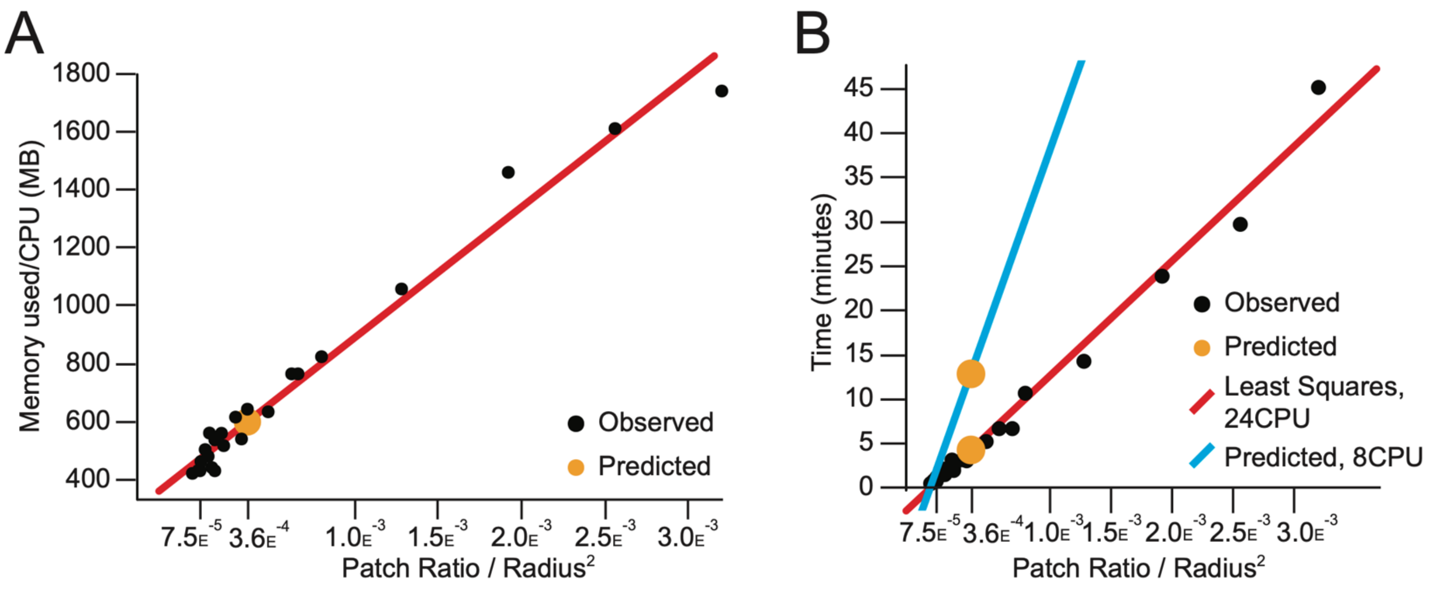


**Supplementary Figure 7. Example computational resource demands of TDAExplore.** Computational resources required for 5-fold cross validation with 25 choices of hyperparameters using PFN1 KO data set (n=68 images). **(A)** Memory usage (maximum resident size) per CPU thread. **(B)** Observed and predicted computation time (wall time) using 24 CPU threads and 8 CPU threads, respectively.

| Dataset | Image resolution (pixels) | Number of images | Images per class |
|---|---|---|---|
| BBBC013 (4) | 640 x 640 | 32 | High concentration: 16<br>Low concentration: 16 |
| BBBC014 (4) | 1360 x 1024 | 32 | High concentration: 16<br>Low concentration: 16 |
| BBBC015 (4) | 1000 x 768 | 12 | High concentration: 6<br>Low concentration: 6 |
| BBBC016 (4) | 512 x 512 | 24 | High concentration: 12<br>Low concentration: 12 |
| Binucleate (5) | 1280 x 1024 | 41 | Binucleated: 20<br>Non-binucleated: 21 |
| Liver “Ad Libium” (5) | 1388 x 1040 | 265 | Female: 115<br>Male: 150 |
| Liver “Calorie Restriction” (5) | 1388 x 1040 | 303 | Female: 153<br>Male: 150 |
| PFN1 KO (1) | 2048 x 2048 | 68 | Control KO: 41<br>PFN1 KO: 27 |
| Cofilin KD (2) | Varies, approximately 2500 x 2500 | 53 | Control KD: 25<br>Cofilin KD: 28 |
| CK666 (1) | 2048 x 2048 | 57 | CK689: 25<br>CK666: 32 |
| Tβ4 KD (3) | 2048 x 2048 | 58 | Control KD: 29<br>Tβ4 KD: 29 |
| FP4-Mito (1) | 2048 x 2048 | 72 | AP4: 36<br>FP4: 36 |
| CCCP (this study) | 2048 x 2048 | 58 | DMSO: 29<br>FCCP: 29 |

**Supplementary Table 1.** Summaries of image datasets used for assessment.

| Dataset | Subject Type | TDAExplore (image) | TDAExplore (patches) |
|---|---|---|---|
| PFN1 KO (1) | Murine CAD cells | 100(0) | 80.9(0.3) |
| Cofilin KD (2) | Murine CAD cells | 90.2(2.1) | 70.1(0.5) |
| CK666 (1) | Murine CAD cells | 90.9(2.3) | 69.7(0.4) |
| Tβ4 KD (3) | Murine CAD cells | 88.1(2.7) | 65.3(0.5) |
| FP4-Mito (1) | Murine CAD cells | 82.3(4.4) | 63.5(1.1) |

**Supplementary Table 2.** TDAExplore classification accuracies for actin cytoskeleton perturbations. Whole image and patch accuracies are reported as percentage of images correctly classified, standard deviation values are listed in parenthesis.

| Dataset | Subject Type | CP-CHARM | M-CNN | TDAExplore (image) | TDAExplore (patches) |
|---|---|---|---|---|---|
| BBBC013 (4) | Human U2OS cells | 99(1) | 100(0) | 100(0) | 93.8(0.5) |
| BBBC014 (4) | MCF-7 cells | 84(3) | 100(14) | 93.6(3.9) | 67.8(1.3) |
| BBBC015 (4) | Human U2OS cells | 99(0.8) | 100(0) | 100(0) | 91.9(0.7) |
| BBBC016 (4) | Human U2OS cells | 81(7) | 100(0) | 100(1.7) | 79.4(1.5) |
| Binucleate (5) | D. Melanogaster cells | 95(2) | - | 100(1.1) | 80.1(0.1) |
| Liver gender (ad libium) (5) | Murine tissue | 98(0.5) | - | 90.9(1.4) | 65.1(1.4) |
| Liver gender (caloric restriction) (5) | Murine tissue | 99(0.1) | - | 89.4(0.8) | 82.4(0.2) |

**Supplementary Table 3.** TDAExplore classification accuracies for third party data sets. Whole image and patch accuracies are reported as percentage of images correctly classified, standard deviation values are in parenthesis. Median accuracy is used for BBBC and IICBU datasets, allowing for comparison with previously reported results from CP-CHARM and M-CNN (6,7).
